# Schizophrenia polygenic risk during typical development reflects multiscale cortical organization

**DOI:** 10.1101/2021.06.13.448243

**Authors:** Matthias Kirschner, Casey Paquola, Budhachandra S. Khundrakpam, Uku Vainik, Neha Bhutani, Benazir Hodzic-Santor, Foivos Georgiadis, Noor B. Al-Sharif, Bratislav Misic, Boris Bernhardt, Alan C. Evans, Alain Dagher

## Abstract

Schizophrenia is widely recognized as a neurodevelopmental disorder. Abnormal cortical development may by revealed using polygenic risk scoring for schizophrenia (PRS-SCZ). We assessed PRS-SCZ and cortical morphometry in typically developing children (3–21 years) using whole genome genotyping and T1-weighted MRI (n=390) from the Pediatric Imaging, Neurocognition and Genetics (PING) cohort. We contextualise the findings using (i) age-matched transcriptomics, (ii) histologically-defined cytoarchitectural types and functionally-defined networks, (iii) case-control differences of schizophrenia and other major psychiatric disorders. Higher PRS-SCZ was associated with greater cortical thickness, which was most prominent in areas with heightened gene expression of dendrites and synapses. PRS-SCZ related increases in vertex-wise cortical thickness were especially focused in the ventral attention network, while koniocortical type cortex (i.e. primary sensory areas) was relatively conserved from PRS-SCZ related differences. The large-scale pattern of cortical thickness increases related to PRS-SCZ mirrored the pattern of cortical thinning in schizophrenia and mood-related psychiatric disorders. Age group models illustrate a possible trajectory from PRS-SCZ associated cortical thickness increases in early childhood towards thinning in late adolescence, which resembles the adult brain phenotype of schizophrenia. Collectively, combining imaging-genetics with multi-scale mapping, our work provides novel insight into how genetic risk for schizophrenia impacts the cortex early in life.

## Introduction

Schizophrenia is a multifaceted and highly heritable psychiatric disorder that is widely recognized to have a neurodevelopmental origin [1–3]. Abnormal brain development likely predates the onset of clinical symptoms, which typically emerge in early adulthood [4]. Genome-wide association studies (GWAS) support this hypothesis by showing schizophrenia-related genes are involved in multiple neurodevelopmental processes [1, 5]. These genes may affect brain development, leading to vulnerability to environmental effects, and have been suggested to contribute to atypical cortical morphology, as previously observed in cohorts with a schizophrenia diagnosis [3].

Childhood and adolescent brain development involves dynamic and complex structural changes that are shaped by genetic and environmental factors [6–10]. Longitudinal neuroimaging studies have consistently reported a global increase in cortical volume, thickness, and surface area that typically peaks in late childhood and is followed by decreases in adolescence [7, 10, 11]. At the same time, regional maturational trajectories are heterochronous, whereby sensory areas mature earlier than transmodal cortex [10, 12, 13], shaping large-scale patterns of cortical differentiation [14].

The neurodevelopmental hypothesis of schizophrenia posits that cortical maturation is perturbed, producing widespread cortical abnormalities [15]. Differences in cortical morphometry are consistently reported across different stages and clinical phenotypes of the schizophrenia spectrum [16–18]. Investigating neurodevelopmental features of schizophrenia requires a departure from classic case-control designs, however. Alternatively, focusing on genetic risk enables us to investigate neuroanatomical correlates in a large population-based cohort of children and adolescents, without interacting disease-related factors (e.g. medication, chronicity). Recent work shows sensitivity of polygenic risk scoring for schizophrenia (PRS-SCZ) to cortical morphometry [19–22], though not necessarily grey matter volume [23–25]. Thus far, studies have centred almost exclusively on adult cohorts. Only one study has investigated adolescents (aged 12–21 years) and noted an association of PRS-SCZ with globally decreased cortical thickness amongst cannabis users [19]. Discerning neurodevelopmental aspects of genetic risk for schizophrenia requires investigation of younger cohorts.

Understanding the relation of genetic risk for schizophrenia to neurodevelopment can be further enhanced by contextualising imaging-derived phenotypes of polygenic risk with maps of cortical organization. At the cellular level, a range of processes associated with healthy cortical development, such as synaptic pruning, dendritic arborization and intracortical myelination, are implicated in the development of schizophrenia and may produce regional cortical disruptions [26–29]. Recent advances in RNA sequencing (RNAseq) of post mortem brain tissue [30] allow discernment of the relative contribution of cell-types to patterns of atypical cortical morphometry [31, 32]. More complex interactions of microstructure and function on regional vulnerability may be captured by the groupings of cortical areas into cytoarchitectural types and functional networks. Indeed, recent studies of schizophrenia [17, 33] and high PRS-SCZ in healthy adults [34] suggest differential sensitivity of histological-defined cytoarchitectural types [35] and functional networks [36]. Finally, population-level effects of schizophrenia and other major psychiatric disorders can be used to illustrate the concordance of genetic risk for schizophrenia with disorder-related neuroanatomical phenotypes. Specifically, it can be tested how the association between genetic risk of schizophrenia and cortical morphometry in children relates to shared and divergent neuroanatomical abnormalities across psychiatric disorders [32]. Taken together, multiple scales of cortical organization can be utilized to provide a comprehensive description of the regional variations of an imaging-derived phenotype, such as genetic risk for schizophrenia.

Here we address the relationship between PRS-SCZ and cortical organization in a large population-based cohort of typically developing children (3 – 21 years) derived from the Pediatric Imaging, Neurocognition and Genetics (PING) study [37]. We hypothesised that higher PRS-SCZ would be associated with atypical cortical morphometry (thickness, surface area and volume). Then, we aimed to better understand the effect of PRS-SCZ on cortical morphometry by comparing the observed spatial patterns to cell-type specific gene expression, cytoarchitectural and functional differentiation, and cortical abnormalities seen in major psychiatric disorders. Finally, we examined age group specific variations of high PRS-SCZ on cortical morphometry across different neurodevelopmental stages

## Material and Methods

### Subjects

Neuroimaging, demographic and genetic data of typically developing children and adolescents were derived from the PING study [37]. The PING dataset is a wide-ranging, publicly shared data resource comprising cross-sectional data from 1493 healthy subjects evaluated at 10 different sites across the United States. Details of the PING dataset are described elsewhere [37].

### Genomic data

Genomic data processing and calculation of polygenic risk scores followed a recent publication from Khundrakpam and colleagues [38]. Specifically, 550,000 single nucleotide polymorphisms (SNPs) were genotyped from saliva samples using the Illumina Human660W-Quad BeadChip. Details on imputation and preprocessing can be found in the supplementary methods. After SNP imputation and preprocessing, 4,673,732 variants were available for calculation of polygenic scores. Participants were filtered to have at least 0.95 loadings to the European genetic ancestry factor (coded as “GAF_europe” in the PING dataset), resulting in 526 participants. To capture/ quantify population structure, the same participants were used to calculate the 10 principal components across the variants, excluding areas in high LD with each other (--indep-pairwise 50 5 0.2) with Plink 2.

The PRS-SCZ was trained using results from latest GWAS on schizophrenia at the time of analysis [39]. The GWAS was filtered for having imputation quality over 90. Polygenic scores were calculated with PRSice 2.30e [40]. Clumping of the data was performed using PRSice default settings (clumping distance = 250kb, threshold r2 = 0.1). To calculate the PRS-SCZ we used the GWAS hits (p< 5×10^*−08^) cut-off criterion. This resulted in 87 variants common to the base and target datasets. (Table S1). The choice of the GWAS hits threshold was made to increase the specificity of observed gene-brain associations for schizophrenia and to minimize the genetic overlap with other psychiatric disorders such as bipolar disorder, which increases with lower PRS significance thresholds by including more SNPs. To illustrate this influence of PRS significance thresholds on the genetic overlap between schizophrenia and bipolar disorder, we calculated PRS for bipolar disorder (PRS-BIP) applying exactly the same processing pipeline (Supplementary Methods). In the present sample, PRS-SCZ and PRS-BIP did not correlate (r=0.063) using the GWAS hits threshold, whereas applying lower significance thresholds, such as p=0.05 and p=0.1, resulted in moderate correlations (r=0.254 and r=0.286, respectively; Figure S1).

### Image acquisition and pre-processing

Details on image acquisition and pre-pre-processing are described elsewhere [37]. The CIVET processing pipeline, (http://www.bic.mni.mcgill.ca/ServicesSoftware/CIVET, page 2.1) [41] was used to compute cortical thickness, surface area and cortical volume measurements at 81,924 regions covering the entire cortex and quality control (QC) was performed by two independent reviewers (see supplementary methods for details). After QC of the total 526 subjects that passed filtering for European genetic ancestry, a total sample of 390 participants remained for all subsequent analyses. Demographic data from the 390 resulting participants from the PING dataset are described in Table 1.

### Statistical analyses

#### Association between PRS-SCZ and cortical morphometry

To identify the association between PRS-SCZ on vertex-wise cortical thickness, surface area and cortical volume, general linear models (GLM) were applied using the SurfStat toolbox (http://www.math.mcgill.ca/keith/surfstat/) [42]. Each cortical feature was modelled as:

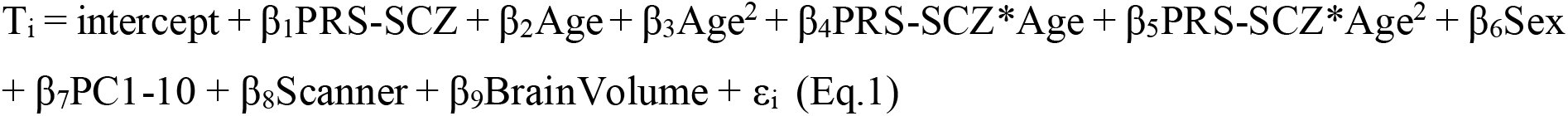

where *i* is a vertex, PRS-SCZ is the polygenic risk score for schizophrenia, *Age* is years at the time of scan, PC10 are the first 10 principal components of genomic data to account for population stratification, *ε* is the residual error, and the intercept and the *β* terms are the fixed effects. Models with quadratic age terms were chosen as they fit the data better than models with only lower degree age terms (Supplementary Methods). For each cortical feature, vertex-wise *t*-statistics of the association with PRS-SCZ (β_1_PRS-SCZ) were mapped onto a standard surface. To assess the significance of PRS-SCZ effects on each of the three different cortical features, whole-brain correction for multiple comparisons using Random field theory (RFT) at cluster-level *p* ≤ .01 [43, 44] was applied. Please note, that only cortical thickness showed a significant association with PRS-SCZ after RFT correction (see Results, Figure 1); all subsequent analyses were thus restricted to cortical thickness only.

#### Cellular composition of the cortex and PRS-SCZ effects on cortical thickness

We evaluated how the observed pattern of PRS-SCZ effects on cortical thickness relates to regional variations in the cellular compositions of the cortex. Given prior evidence [45] and histological validation (Supplementary Methods), we focused on components of the neuropil, namely glial cell processes, axons, dendritic trees, neuron-to-neuron synapses i.e. in cortical tissue other than cell bodies or blood vessels. Neuropil-related gene expression was calculated by combining tissue-level RNAseq with single-cell RNAseq for cell-types [30] and gene ontologies for neuron-compartments [46, 47]. Tissue-level RNAseq provided expression levels of 60,155 genes in 11 neocortical areas. The areas were cytoarchitecturally defined in each specimen, supporting precise mapping and comparison across individuals. Crucially, we selected 12 brain specimens that were age-matched to the PING imaging cohort (3-21 years), because gene expression differs substantially between children and adults [48]. Single-cell RNAseq provided specificity scores for each gene to glial cell types. For each type, we weighted the genes by the specificity score, then calculated the average across genes, across specimens and within area. For each neuron-compartment, we defined a list of marker genes using The Gene Ontology knowledgebase, then calculated the average expression of marker genes in each area and specimen. The annotated terms used were “neuron_to_neuron_synapse”, “dendritic_tree” and “main_axon”. Next, we mapped the 11 areas to the cortical surface and extracted area-average PRS-SCZ effects on cortical thickness. The cortical areas were visually matched to nearest parcel in a 200 parcel decomposition of the Desikan-Killany, as performed in previous work [49]. Finally, we tested the spatial similarity of cell-type specific gene expression with PRS-SCZ effects on cortical thickness using product-moment correlations. Statistical significance was determined relative to random reassignment permutation tests (10,000 repetitions).

**Figure 1:**
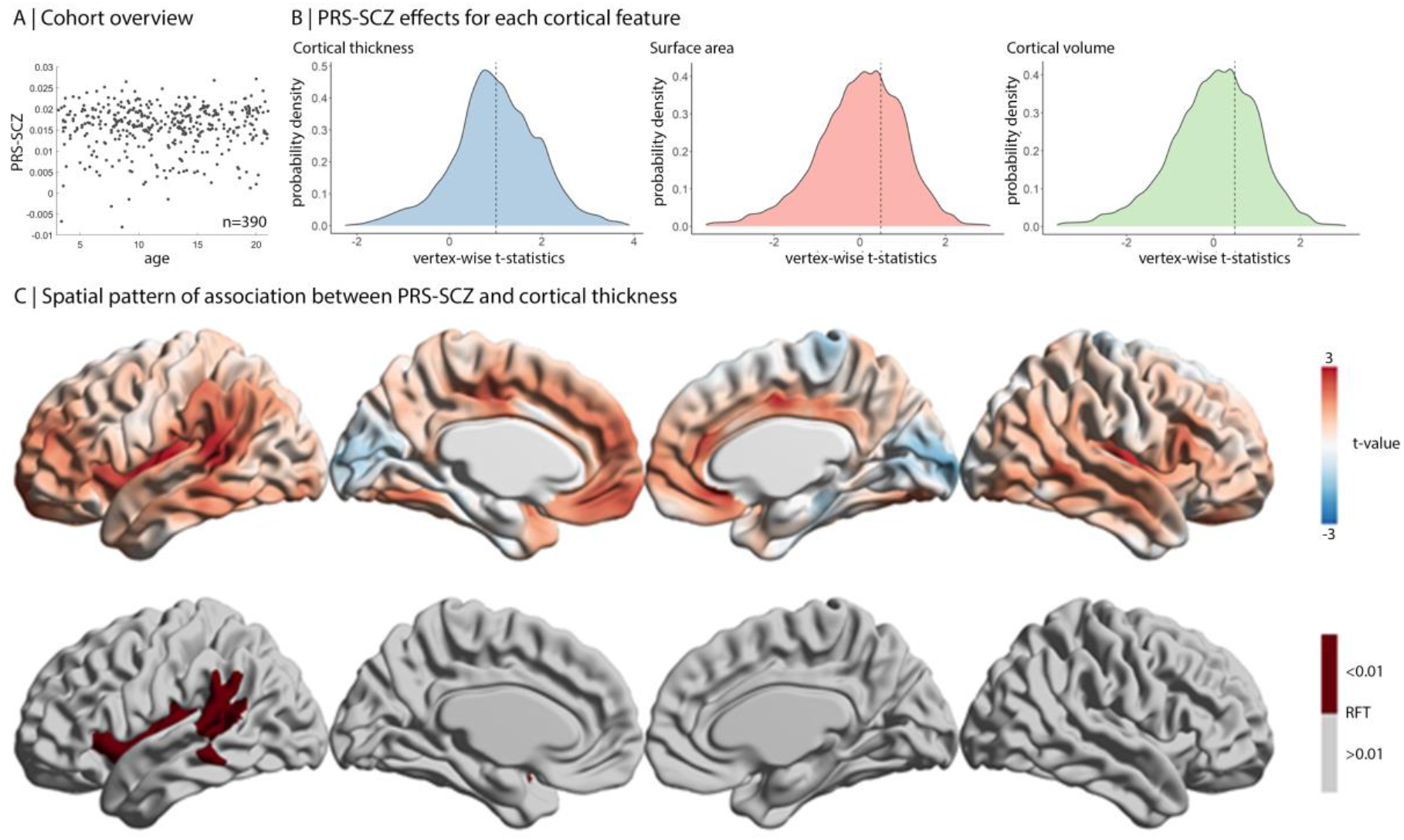
Association of PRS-SCZ with cortical morphometry. **A)** Scatterplot shows variability of PRS-SCZ scores across age range within the selected cohort. **B)** Probability distributions show the variation in vertex-wise t-values of the association of PRS-SCZ with each cortical feature. Dashed lines represent the mean values of each cortical feature. Only cortical thickness was significantly shifted from 0. **C)** Unthresholded (top) and thresholded (bottom) maps after RFT correction (P<0.01) show the association of PRS-SCZ with cortical thickness. Unthresholded maps for surface area and cortical volume are Supplementary Figures 2-3.

#### Aggregation of PRS-SCZ effects on cortical thickness by cytoarchitectural type and functional network

We contextualized the PRS-SCZ effects on cortical thickness by cytoarchitectural types and intrinsic functional networks. Cytoarchitectural types were assigned to Von Economo areas [50, 51], based on a recent re-analysis of Von Economo micrographs [35]. Cortical types synopsize degree of granularity, from high laminar elaboration in koniocortical areas, six identifiable layers in eulaminate III-I, poorly differentiated layers in dysgranular and absent layers in agranular.

Functional networks were defined based on the Yeo atlas [36]. The atlas reflects clustering of cortical vertices according to similarity in resting state functional connectivity profiles, acquired in 1000 healthy young adults. We assessed whether the PRS-SCZ effects on cortical thickness were stronger or weaker within each class, relative to spin permutations [52, 53]. Specifically, we calculated the median PRS-SCZ effect for each cytoarchitectural type and each functional network, then compared the median scores to a null model, which was constructed by randomly rotating the atlases across the cortical surface and re-evaluating the median scores (10,000 permutations). The medial wall was assigned as a NaN and not included in each the permuted correlation [54]. Statistical significance was deemed where p_spin_<0.025 (two-tailed test).

#### Pattern similarity between PRS-SCZ effects and cortical abnormalities in major psychiatric disorders

We assessed whether cortical thickness differences of PRS-SCZ relate to patterns of cortical thickness abnormalities observed in major psychiatric disorders including schizophrenia, bipolar disorder, major depressive disorder (MDD), attention deficit hyperactivity disorder (ADHD), autism spectrum disorder (ASD), obsessive compulsive disorder (OCD). To this end, the PRS-SCZ related t-statistic map was parcellated into 64 Desikan-Killiany (DK) atlas regions [55] and then correlated with the corresponding Cohen’s d maps derived from recently published meta-analyses by the ENIGMA schizophrenia [15], bipolar disorder [56], major depressive disorder working groups [57], attention deficit hyperactivity disorder [58], obsessive compulsive disorder [59] and autism spectrum disorder [60] implemented in the ENIGMA toolbox [61]. Specifically, spatial pattern similarity of cortical DK maps was examined using product-moment correlations. Statistical significance and correction of spatial autocorrelation were assessed with the spin permutation tests (10,000 repetitions) [52, 53] implemented in the ENIGMA toolbox [61]. The medial wall was assigned as a NaN and not included in each the permuted correlation [54]. Statistical significance was deemed where p_spin_<0.025 (two-tailed test) and false discovery rate (FDR) (pFDR < 0.05) was applied to control for multiple comparisons (n=6).

#### Age group effects of PRS-SCZ on cortical thickness

We examined the distinctiveness of PRS-SCZ related cortical thickness differences in different neurodevelopmental stages by dividing the sample into three age groups; early childhood (3-9years, n=145), early adolescence (10-15years, n=155) and late adolescence (16-21years, n=116) [62]. The PRS-SCZ effect on cortical thickness was evaluated within each group using the abovementioned GLM (Eq.1), however, the age term was centred to the mean of the group to focus on the effect within the specified developmental stage [8]. Note that there was no correlation between age and PRS-SCZ scores (r=0.058, Figure 1A). We specifically examined whether the association of the PRS-SCZ effect with the adult phenotype of SCZ changed across age groups. To do so, we compared the product-moment correlation coefficients between maps [63].

## Results

### Polygenic risk for SCZ is associated with greater cortical thickness

To test the association between PRS-SCZ and cortical morphometry in typically developing children, we used T1-weighted MRI and whole genome genotyping (n=390) from the PING cohort (3–21 years, mean±sd = 12.1±4.7 years, 46% female) (**Table S2**). Vertex-wise GLMs related cortical thickness with PRS-SCZ, controlling for age, sex, the first 10 principal components of genetic variants (to account for population stratification), scanner, and total brain volume. We found that higher PRS-SCZ was significantly associated with greater cortical thickness (Random Field Theory (RFT) corrected, P < 0.01) but not surface area or cortical volume (**Figure 1B, Figures S2-3**). Overall, the unthresholded t-statistic map revealed that higher PRS-SCZ was associated with widespread increases in cortical thickness in association cortex, but reduced cortical thickness in sensory areas (**Figure 1C top**). Higher PRS-SCZ was associated with significantly thicker cortex in the left insula, left superior temporal gyrus and left inferior parietal lobule (**Figure 1C bottom**, RFT corrected, P<0.01). These results suggest a significant effect of PRS-SCZ on cortical thickness but not surface area or cortical volume in typically developing children. As such, subsequent analyses are restricted to cortical thickness. In a next step, we sought to examine how the PRS-SCZ related cortical thickness increase in typically developing children relate to different levels of cortical organization including 1) cell type specific gene expression 2) cytoarchitectural and functional systems and 3) cortical pattern of case-control differences from schizophrenia and other major psychiatric disorders.

### Alignment with cell-type specific gene expression

Histological examinations have reported a null or minimal relationship between cortical thickness and neuron number in healthy brain samples [64–66]. Instead, regional variations in cortical thickness show a strong association with neuropil [66], the portion of cortical tissue that excludes cell bodies or blood vessels [67]. We examined whether cortical thickness differences related to PRS-SCZ mirrored regional variations in the neuropil composition of the cortex, in order to generate hypotheses on the neuropil components affected by PRS-SCZ. To this end, we first validated the relationship between cortical thickness and neuropil using tabular data and photomicrographs of Nissl stains [r=0.49, p_spin_=.018, **Figure 2A**, [50, 68]]. In contrast, neuronal density was not correlated with histologically-defined cortical thickness (r=0.08 p_spin_=.678), which aligns with previous work [65, 66]. Then, we estimated regional variations in neuropil-related gene expression, based on six cellular components (astrocytes, microglia, oligodendrocytes, axons, dendritic trees, neuron-to-neuron synapses) by combining tissue-level RNAseq with single-cell RNAseq for cell-types [30] and gene ontologies for neuron-compartments [46, 47] (**Figure 2B**). Correlating PRS-SCZ effects on cortical thickness with neuropil-related gene expression, we found that PRS-SCZ effects are significantly associated with gene expression for dendritic trees (r=.755, p_perm_=.006), synapses (r=.618, p_perm_=.005), and at a trend-level with axons (r=.481, p_perm_=.069) (**Figure 2C**). In contrast, no significant correlation was observed with gene expression related to glial components of neuropil (**Figure 2C**). Together, these analyses suggest greater cortical thickness with higher PRS-SCZ is observed in areas with greater dendritic and synaptic density.

**Figure 2:**
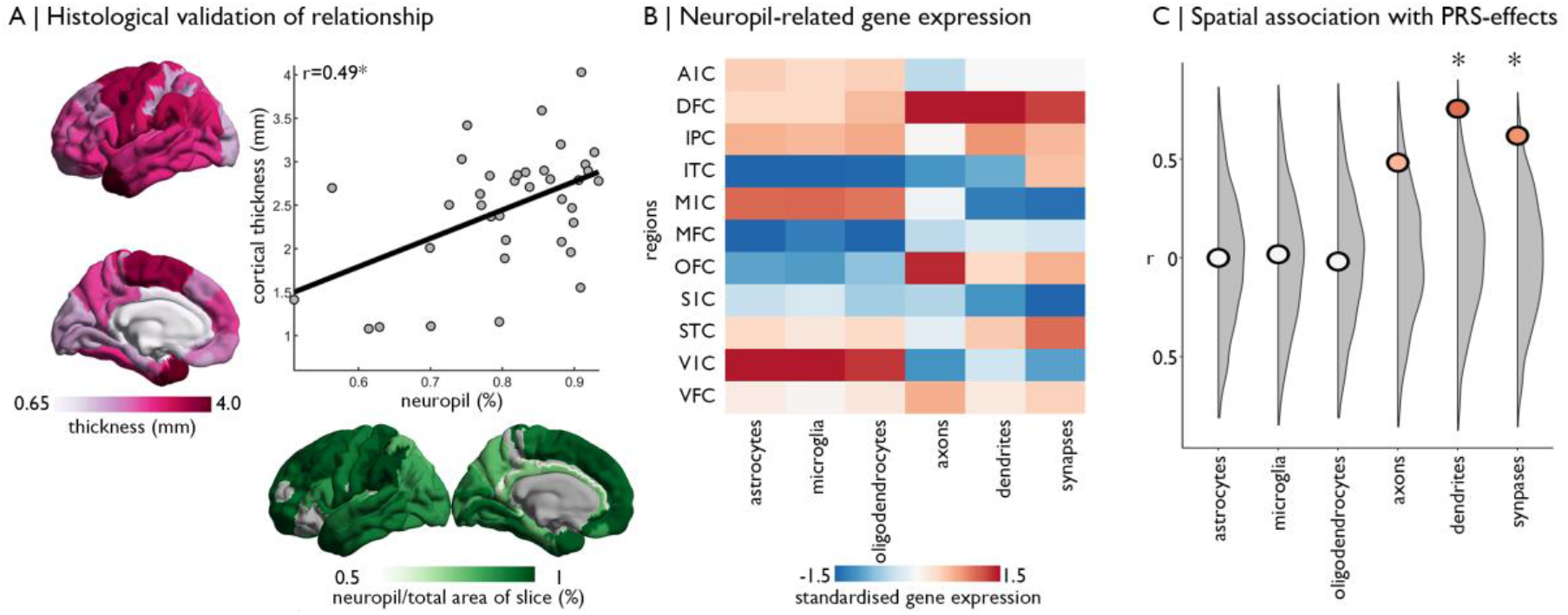
Decoding spatial patterns of PRS-SCZ on cortical thickness. **A)** Correlation of histological measurements of cortical thickness and neuropil from Von Economo & Koskinas [50]. Cortical thickness is shown in pink and on the y-axis. Neuropil is shown in green and on the x-axis. **B)** Gene expression varies across glial and neuron-related compartments of neuropil in eleven cortical regions. **C)** Correlation of neuropil-related gene expression with the PRS-SCZ effects showed significant association with dendrites and synapses, compared with null distributions from permutation testing (grey). **Abbreviations**. A1C: primary auditory cortex. DFC: dorsal frontal cortex. IPC: inferior parietal cortex. ITC: inferior temporal cortex. M1C: primary motor cortex. MFC: medial frontal cortex. OFC: orbital frontal cortex. S1C: primary somatosensory cortex. STC: superior temporal cortex. V1C: primary visual cortex. VFC: ventral frontal cortex.

**Figure 3:**
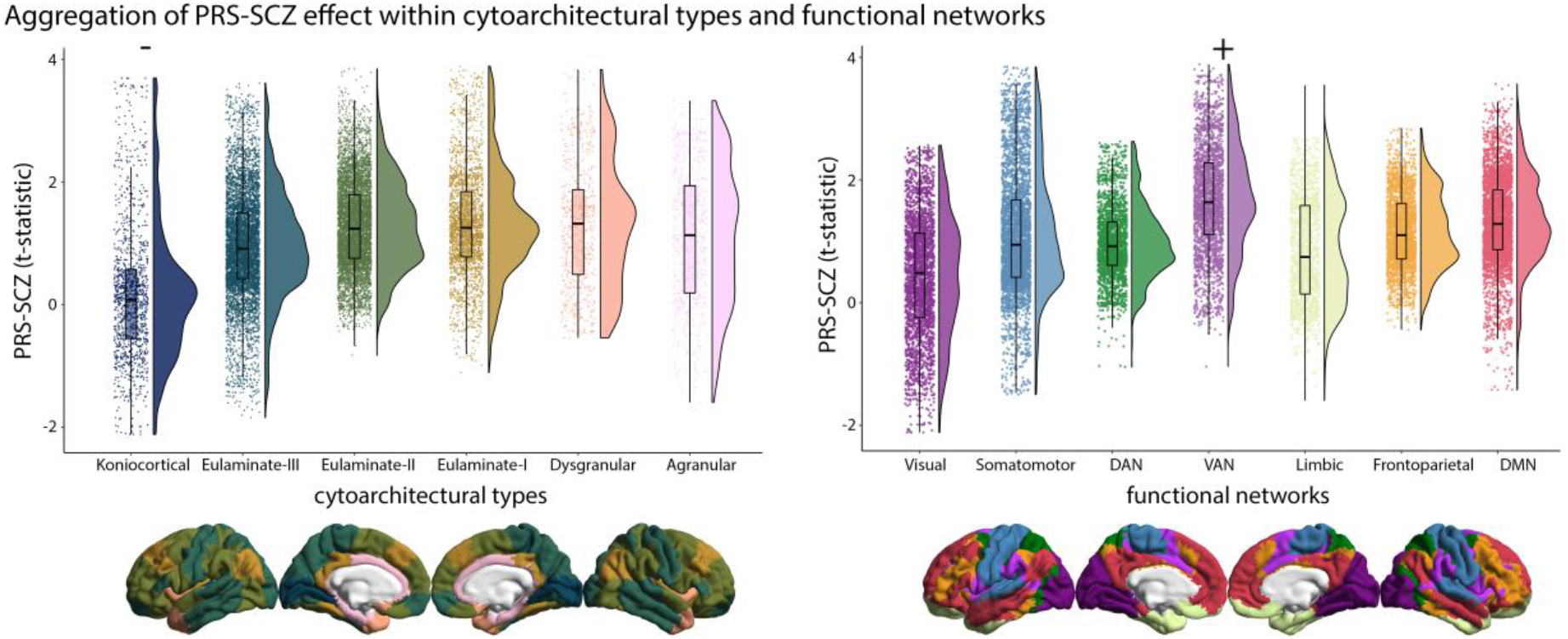
Aggregation of PRS-SCZ effect within cytoarchitectural types and functional networks. Raincloud plots show the distribution of the PRS-SCZ effect on cortical thickness stratified by cytoarchitectural type [35] and functional network [36]. Relative to spin permutation null models, the koniocortical cortical type encompassed significantly lower t-statistics, whereas the VAN encompassed significantly higher t-statistics. DAN: dorsal attention network. VAN: ventral attention network. DMN: default mode network.

### Contextualisation by cytoarchitectural types and functional networks

Next, we aimed to determine whether the PRS-SCZ is preferentially associated with certain cytoarchitectural types or functional networks. Based on established atlases of cytoarchitectural and functional differentiation [35, 36], we found that the PRS-SCZ effect was stronger, compared to null models, in the ventral attention network (median±SD=1.64±0.88, p_spin_=0.004). Conversely, PRS-SCZ effect was weaker or more negative, compared to null models, in the koniocortical type (i.e. primary sensory areas) (median±SD=0.08±1.10, p_spin_=0.008).

### Cortical thickness signatures of PRS-SCZ and major psychiatric disorders

We assessed whether PRS-SCZ effects on cortical thickness relate to abnormal cortical thickness patterns observed in case-control meta-analyses (Cohen’s d-maps) of schizophrenia and other psychiatric illnesses. The PRS-SCZ related cortical thickness increase showed a negative correlation with schizophrenia-related cortical abnormalities schizophrenia (r=−.326, p_spin_=.0022). In addition, we found similar negative correlations to cortical abnormalities in BD (r=−.466, p_spin_<.001), MDD (r=−.538, p_spin_<.001) and ADHD (r=−.430, p_spin_<.001) but not OCD or ASD (**Figure 4**). Altogether, cortical regions showing PRS-SCZ related greater thickness are those with the strongest thinning across disease maps of schizophrenia and genetically related affective disorders (e.g. BD, MDD) and ADHD.

**Figure 4:**
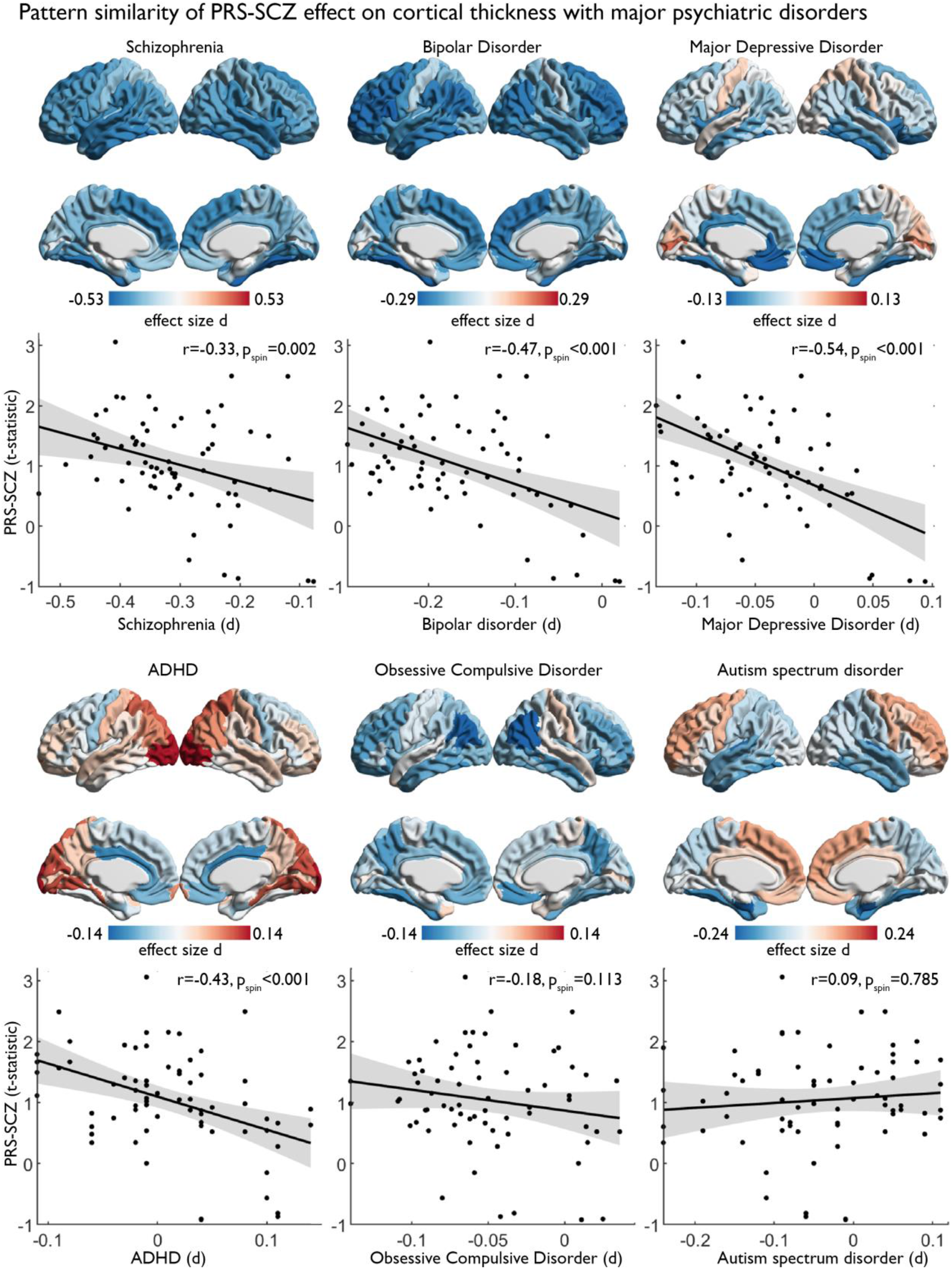
Pattern similarity of PRS-SCZ on cortical thickness with major psychiatric disorders. Cortical surfaces show the effect size of schizophrenia, bipolar disorder and major depressive disorder, attention deficit hyperactivity disorder, obsessive compulsive disorder and autism spectrum disorder diagnosis on cortical thickness from ENIGMA meta-analyses of each disorder [61]. Statistical significance was deemed where pspin<0.025 (two-tailed test) and false discovery rate (FDR) (pFDR < 0.05) was applied to control for multiple comparisons (n=6). Scatterplots show the correlation of each map with PRS-SCZ effect on cortical thickness.

### Age group effects of PRS-SCZ

Schizophrenia-related genes are implicated in neurodevelopmental processes and as such the effect of PRS-SCZ on cortical thickness likely varies with age. Although the cross-sectional nature of this cohort prohibits mapping individual trajectories of cortical development, we sought to approximate developmental variation in the effect of PRS-SCZ by estimating age group effects of early childhood (3-9 yrs), early adolescence (10-15 yrs), and late adolescence (15-21) in the cohort. Higher PRS-SCZ was associated with greater cortical thickness in early childhood, however the pattern differs in the older age groups (**Figure 5**). We detected a significant difference in the correlation coefficients between early childhood and late adolescence (z=2.84, p=0.002), as well as early adolescence and late adolescence (z=1.84, p=0.033). Furthermore, PRS-SCZ related cortical thickness increase in early childhood correlated negatively with schizophrenia-related cortical abnormalities, whereas PRS-SCZ related cortical thinning in late adolescents correlated positively (**Figure 5**). To further inspect the age-related change in the PRS-SCZ effect on cortical thickness, we repeated the analysis using the entire cohort and iteratively shifted the age-centring from 3-21 in 1yr intervals. Higher PRS-SCZ was associated with greater cortical thickness in the 3-6yr age-centred models (Figure S4) closely resembling the results from the main analysis (**Figure 1C**).

**Figure 5:**
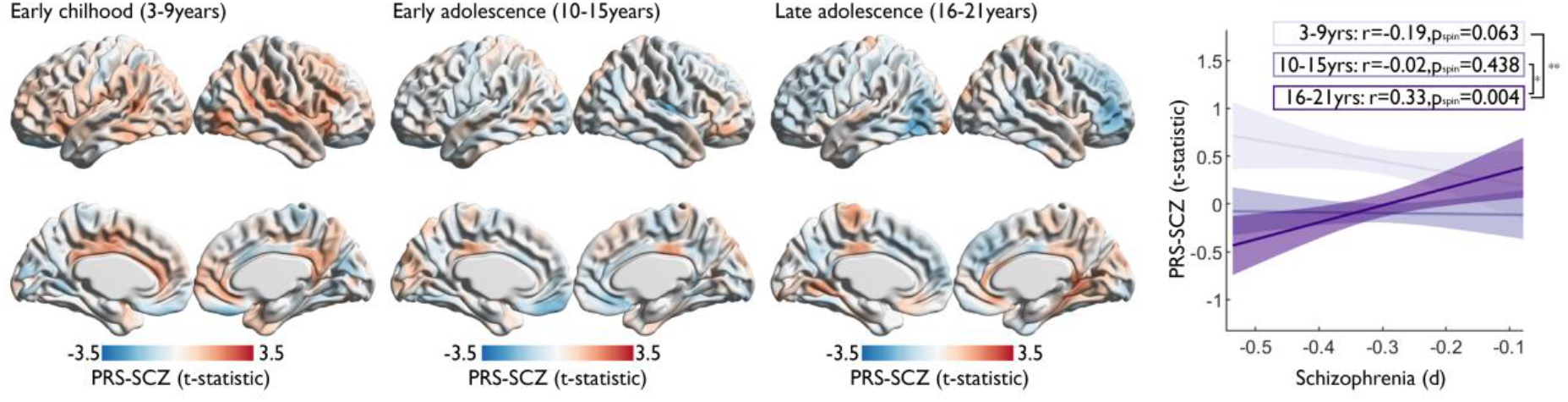
Age group based PRS-SCZ effect on cortical thickness. Cortical surfaces show unthresholed maps of PRS-SCZ effect on cortical thickness within age groups. Line plot shows how the relationship (linear regression with 95% confidence intervals) of PRS-SCZ with the SCZ-related pattern of abnormalities (Figure 4) changes from negative to positive across the age groups. Significant difference in correlation coefficients is shown by * for p<0.05 and ** for p<0.01.

## Discussion

Combining imaging-genetics with multi-scale mapping, we characterised the relation of PRS-SCZ on cortical morphometry across different scales of cortical organization. We found that higher PRS-SCZ was associated with greater cortical thickness in typically developing children, while surface area and cortical volume showed only subtle associations with PRS-SCZ. We further provide evidence that PRS-SCZ preferentially affects areas with heightened expression of dendrites and synapses and that the PRS-SCZ related cortical differences accumulate in cytoarchitecturally and functionally defined cortical systems. We also find that the PRS-SCZ related cortical pattern mirrors cortical thinning related to schizophrenia and other major psychiatric disorders. Finally, age group models suggest a potential trajectory from PRS-SCZ associated cortical thickness increase in early childhood towards thinning in late adolescence spatially resembling the adult brain phenotype of schizophrenia.

Our cell-type specific gene expression approach enabled cross-modal exploration of the relationship of genetic risk for schizophrenia with expression levels of neurons and glia. Our findings support and extend upon post-mortem analyses, which demonstrate abnormal dendritic and synaptic density in individuals with schizophrenia (see [70] for a recent meta-analysis). Another recent study showed that differences in cortical thickness across multiple psychiatric disorders (including schizophrenia) are associated with pyramidal-cell gene expression, a gene set enriched for biological processes of dendrites (e.g. dendritic arborization and branching) [71] as well as synaptic function [32]. The present findings extend this work by demonstrating a relationship between gene expression of dendrites and synapses with PRS-SCZ related cortical differences during neurodevelopment.

System-specific contextualization of the PRS-SCZ effects on cortical thickness revealed that the ventral attention network was preferentially sensitive to PRS-SCZ, while koniocortical type cortex (e.g. primary sensory areas) was mostly spared from its influence. This cortical thickness pattern of PRS-SCZ closely mirrors recent observations in patients with schizophrenia showing stronger brain abnormalities in the ventral attention network, while primary cortex, as defined by von Economo, was relatively spared [33]. Altogether, these findings demonstrate that system-specific differentiations of PRS-SCZ related cortical thickness differences during neurodevelopment reflect cortical abnormalities of schizophrenia suggesting some neuroanatomical continuity between polygenic risk and clinical phenotype.

Longitudinal data and case-control meta-analysis have shown that the development of psychosis in high-risk adolescents is associated with progressive loss of cortical thickness in several areas of the association cortex [18, 72]. Of note, the areas implicated in these studies overlap considerably with those showing increased cortical thickness in early childhood and more pronounced cortical thinning in adolescents with higher PRS-SCZ in the current study. We further observed that the pattern of PRS-SCZ related cortical thickening was associated with areas of cortical thinning in schizophrenia, BD, MDD, ADHD. This transdiagnostic overlap notably mirrors the genetic and phenotypic correlation between these disorders [32, 73]. While we do not know the cause of increased cortical thickness in our sample, converging evidence supports the idea that reduced cortical thickness in adults with schizophrenia results from loss of neuropil, and specifically synapses. For example, post mortem studies in schizophrenia demonstrate synaptic loss [70]; many genes implicated in schizophrenia are associated with synapses or synaptic pruning [5, 74, 75]; regional variations in cortical thickness correlate with neuropil ([66]. It is conceivable that the PRS-SCZ is associated with delayed pruning and an excess of synapses for age, which in turn may render the affected brain regions vulnerable to catastrophic synaptic loss during the emergence of psychosis.

The association of PRS-SCZ with greater cortical thickness in early childhood raises the question of how the genetic risk of schizophrenia contributes to abnormal developmental trajectories. Given that the transmodal areas identified in the present analysis, such as the insula, exhibit modest cortical thinning from 3-21 years [76–78], our results align with either an amplified trajectory (i.e.: higher peak, steeper decline) and/or delayed cortical thinning in early childhood. Related to the complexity and heterochronicity of cortical maturation during childhood and adolescents [76], polygenic disorders can involve multiple types of abnormal trajectories, occurring simultaneously or sequentially [79, 80]. Amplified or delayed trajectory of transmodal area morphometry may represent a core motif of cortical development in children with high PRS-SCZ. The PRS-SCZ includes multiple genetic factors, however, and their individual variation may produce heterogeneity in cortical development within high PRS-SCZ individuals.

The present study should be interpreted in the light of the cross-sectional nature of the dataset that limits the ability to map individual longitudinal trajectories. The convergence of the observed findings with gene expression and neuroanatomical studies of schizophrenia support multiscale continuity between polygenic risk and clinical phenotype of schizophrenia. However, given the low familial risk and relative absence of other biological or environmental risk factors for schizophrenia in the study cohort, interaction between PRS-SCZ and other biological and environmental risk factors could not directly be assessed and warrants further investigation. Future research could therefore be enhanced by larger datasets with longitudinal designs and longer follow-up to determine which individuals will develop psychosis or other mental disorders. Finally, the observed PRS-SCZ related cortical thickness increase in early childhood (age 3 to 9) highlights the need for large-scale initiatives targeting this age range.

## Conclusions

The present study provides novel evidence on the cellular basis and developmental trajectory of cortical thickness differences related to genetic risk for schizophrenia that may help to refine the neurodevelopmental hypothesis of schizophrenia. More generally, the present work illustrates how maps of cortical organization can enrich descriptions of imaging-derived phenotypes related to genetic risk for mental illnesses. Altogether, this integrative framework combining imaging-genetics and multi-scale mapping could advance our understanding of the complex associations between individual genetic profiles and cortical organization across multiple psychiatric and neurological conditions.

## Acknowledgments

MK acknowledges funding from the Swiss National Science Foundation (P2SKP3_178175). AD is supported by the Canadian Institutes of Health Research Foundation Scheme. NBA is supported by grants from Brain Canada (238990, 243030), CFREF/HBHL Innovative Ideas (247613), Coutu Research Fund (241177), CFREF/HBHL Discovery (247712).

## Data Availability Statement

All neuroimaging, demographic and genetic data are available from the Pediatric Imaging, Neurocognition, and Genetics (PING) study available at (https://nda.nih.gov/). Tissue-level RNAseq with single-cell RNAseq for cell-types and gene ontologies for neuron-compartments are available at http://development.psychencode.org/. Effect size maps for major psychiatric disorders are available in the ENIGMA toolbox (https://enigma-toolbox.readthedocs.io/en/latest/).

## References

1. Birnbaum R, Weinberger DR. Genetic insights into the neurodevelopmental origins of schizophrenia. Nat Rev Neurosci. 2017;18:727–740.

2. Insel TR. Rethinking schizophrenia. Nature. 2010;468:187–193.

3. Rapoport JL, Giedd JN, Gogtay N. Neurodevelopmental model of schizophrenia: update 2012. Mol Psychiatry. 2012;17:1228–1238.

4. Kessler RC, Angermeyer M, Anthony JC, DE Graaf R, Demyttenaere K, Gasquet I, et al. Lifetime prevalence and age-of-onset distributions of mental disorders in the World Health Organization’s World Mental Health Survey Initiative. World Psychiatry. 2007;6:168–176.

5. Schizophrenia Working Group of the Psychiatric Genomics Consortium, Ripke S, Neale BM, Corvin A, Walters JTR, Farh K-H, et al. Biological insights from 108 schizophrenia-associated genetic loci. Nature. 2014;511:421–427.

6. Shaw P, Greenstein D, Lerch J, Clasen L, Lenroot R, Gogtay N, et al. Intellectual ability and cortical development in children and adolescents. Nature. 2006;440:676–679.

7. Raznahan A, Shaw P, Lalonde F, Stockman M, Wallace GL, Greenstein D, et al. How Does Your Cortex Grow? J Neurosci. 2011;31:7174–7177.

8. Khundrakpam BS, Lewis JD, Kostopoulos P, Carbonell F, Evans AC. Cortical Thickness Abnormalities in Autism Spectrum Disorders Through Late Childhood, Adolescence, and Adulthood: A Large-Scale MRI Study. Cereb Cortex. 2017;27:1721–1731.

9. Park B, Bethlehem RA, Paquola C, Larivière S, Rodríguez-Cruces R, Vos de Wael R, et al. An expanding manifold in transmodal regions characterizes adolescent reconfiguration of structural connectome organization. ELife. 2021;10:e64694.

10. Sydnor VJ, Larsen B, Bassett DS, Alexander-Bloch A, Fair DA, Liston C, et al. Neurodevelopment of the association cortices: Patterns, mechanisms, and implications for psychopathology. Neuron. 2021;109:2820–2846.

11. Tamnes CK, Herting MM, Goddings A-L, Meuwese R, Blakemore S-J, Dahl RE, et al. Development of the Cerebral Cortex across Adolescence: A Multisample Study of Inter-Related Longitudinal Changes in Cortical Volume, Surface Area, and Thickness. J Neurosci. 2017;37:3402–3412.

12. Giedd JN, Blumenthal J, Jeffries NO, Castellanos FX, Liu H, Zijdenbos A, et al. Brain development during childhood and adolescence: a longitudinal MRI study. Nature Neuroscience. 1999;2:861–863.

13. Gogtay N, Giedd JN, Lusk L, Hayashi KM, Greenstein D, Vaituzis AC, et al. Dynamic mapping of human cortical development during childhood through early adulthood. PNAS. 2004;101:8174–8179.

14. Paquola C, Bethlehem RA, Seidlitz J, Wagstyl K, Romero-Garcia R, Whitaker KJ, et al. Shifts in myeloarchitecture characterise adolescent development of cortical gradients. ELife. 2019;8:e50482.

15. van Erp TGM, Walton E, Hibar DP, Schmaal L, Jiang W, Glahn DC, et al. Cortical Brain Abnormalities in 4474 Individuals With Schizophrenia and 5098 Control Subjects via the Enhancing Neuro Imaging Genetics Through Meta Analysis (ENIGMA) Consortium. Biol Psychiatry. 2018;84:644–654.

16. Wannan CMJ, Cropley VL, Chakravarty MM, Bousman C, Ganella EP, Bruggemann JM, et al. Evidence for Network-Based Cortical Thickness Reductions in Schizophrenia. Am J Psychiatry. 2019;176:552–563.

17. Kirschner M, Shafiei G, Markello RD, Makowski C, Talpalaru A, Hodzic-Santor B, et al. Latent Clinical-Anatomical Dimensions of Schizophrenia. Schizophr Bull. 2020. 3 August 2020. https://doi.org/10.1093/schbul/sbaa097.

18. ENIGMA Clinical High Risk for Psychosis Working Group, Jalbrzikowski M, Hayes RA, Wood SJ, Nordholm D, Zhou JH, et al. Association of Structural Magnetic Resonance Imaging Measures With Psychosis Onset in Individuals at Clinical High Risk for Developing Psychosis: An ENIGMA Working Group Mega-analysis. JAMA Psychiatry. 2021. 5 May 2021. https://doi.org/10.1001/jamapsychiatry.2021.0638.

19. French L, Gray C, Leonard G, Perron M, Pike GB, Richer L, et al. Early Cannabis Use, Polygenic Risk Score for Schizophrenia, and Brain Maturation in Adolescence. JAMA Psychiatry. 2015;72:1002–1011.

20. Liu B, Zhang X, Cui Y, Qin W, Tao Y, Li J, et al. Polygenic Risk for Schizophrenia Influences Cortical Gyrification in 2 Independent General Populations. Schizophr Bull. 2017;43:673–680.

21. Neilson E, Bois C, Gibson J, Duff B, Watson A, Roberts N, et al. Effects of environmental risks and polygenic loading for schizophrenia on cortical thickness. Schizophr Res. 2017;184:128–136.

22. Neilson E, Shen X, Cox SR, Clarke T-K, Wigmore EM, Gibson J, et al. Impact of Polygenic Risk for Schizophrenia on Cortical Structure in UK Biobank. Biological Psychiatry. 2019. 22 April 2019. https://doi.org/10.1016/j.biopsych.2019.04.013.

23. Van der Auwera S, Wittfeld K, Homuth G, Teumer A, Hegenscheid K, Grabe HJ. No association between polygenic risk for schizophrenia and brain volume in the general population. Biol Psychiatry. 2015;78:e41–42.

24. Auwera SV der, Wittfeld K, Shumskaya E, Bralten J, Zwiers MP, Onnink AMH, et al. Predicting brain structure in population-based samples with biologically informed genetic scores for schizophrenia. American Journal of Medical Genetics Part B: Neuropsychiatric Genetics. 2017;174:324–332.

25. Reus LM, Shen X, Gibson J, Wigmore E, Ligthart L, Adams MJ, et al. Association of polygenic risk for major psychiatric illness with subcortical volumes and white matter integrity in UK Biobank. Sci Rep. 2017;7.

26. Feinberg I. Schizophrenia: Caused by a fault in programmed synaptic elimination during adolescence? Journal of Psychiatric Research. 1982;17:319–334.

27. Sellgren CM, Gracias J, Watmuff B, Biag JD, Thanos JM, Whittredge PB, et al. Increased synapse elimination by microglia in schizophrenia patient-derived models of synaptic pruning. Nature Neuroscience. 2019;22:374–385.

28. Sprooten E, O’Halloran R, Dinse J, Lee WH, Moser DA, Doucet GE, et al. Depth-dependent intracortical myelin organization in the living human brain determined by in vivo ultra-high field magnetic resonance imaging. NeuroImage. 2019;185:27–34.

29. Wei W, Zhang Y, Li Y, Meng Y, Li M, Wang Q, et al. Depth-dependent abnormal cortical myelination in first-episode treatment-naïve schizophrenia. Human Brain Mapping. 2020;41:2782–2793.

30. Zhu Y, Sousa AMM, Gao T, Skarica M, Li M, Santpere G, et al. Spatiotemporal transcriptomic divergence across human and macaque brain development. Science. 2018;362.

31. Seidlitz J, Nadig A, Liu S, Bethlehem RAI, Vértes PE, Morgan SE, et al. Transcriptomic and cellular decoding of regional brain vulnerability to neurogenetic disorders. Nature Communications. 2020;11:3358.

32. Writing Committee for the Attention-Deficit/Hyperactivity Disorder, Autism Spectrum Disorder, Bipolar Disorder, Major Depressive Disorder, Obsessive-Compulsive Disorder, and Schizophrenia ENIGMA Working Groups. Virtual Histology of Cortical Thickness and Shared Neurobiology in 6 Psychiatric Disorders. JAMA Psychiatry. 2021;78:47–63.

33. Shafiei G, Markello RD, Makowski C, Talpalaru A, Kirschner M, Devenyi GA, et al. Spatial Patterning of Tissue Volume Loss in Schizophrenia Reflects Brain Network Architecture. Biological Psychiatry. 2020;87:727–735.

34. Cao H, Zhou H, Cannon TD. Functional connectome-wide associations of schizophrenia polygenic risk. Mol Psychiatry. 2020:1–9.

35. García-Cabezas MÁ, Hacker JL, Zikopoulos B. A Protocol for Cortical Type Analysis of the Human Neocortex Applied on Histological Samples, the Atlas of Von Economo and Koskinas, and Magnetic Resonance Imaging. Front Neuroanat. 2020;14.

36. Yeo BTT, Krienen FM, Sepulcre J, Sabuncu MR, Lashkari D, Hollinshead M, et al. The organization of the human cerebral cortex estimated by intrinsic functional connectivity. J Neurophysiol. 2011;106:1125–1165.

37. Jernigan TL, Brown TT, Hagler DJ, Akshoomoff N, Bartsch H, Newman E, et al. The Pediatric Imaging, Neurocognition, and Genetics (PING) Data Repository. NeuroImage. 2016;124:1149–1154.

38. Khundrakpam B, Vainik U, Gong J, Al-Sharif N, Bhutani N, Kiar G, et al. Neural correlates of polygenic risk score for autism spectrum disorders in general population. Brain Commun. 2020;2.

39. Ruderfer DM, Ripke S, McQuillin A, Boocock J, Stahl EA, Pavlides JMW, et al. Genomic Dissection of Bipolar Disorder and Schizophrenia, Including 28 Subphenotypes. Cell. 2018;173:1705–1715.e16.

40. Choi SW, O’Reilly PF. PRSice-2: Polygenic Risk Score software for biobank-scale data. Gigascience. 2019;8.

41. Ad-Dab’bagh Y, Einarson D, Lyttelton O, Muehlboeck J-S, Mok K, Ivanov O, et al. The CIVET Image-Processing Environment: A Fully Automated Comprehensive Pipeline for Anatomical Neuroimaging Research 2006. p. 1.

42. Worsley K, Taylor J, Carbonell F, Chung M, Duerden E, Bernhardt B, et al. SurfStat: A Matlab toolbox for the statistical analysis of univariate and multivariate surface and volumetric data using linear mixed effects models and random field theory. NeuroImage. 2009;47:S102.

43. Hayasaka S, Phan KL, Liberzon I, Worsley KJ, Nichols TE. Nonstationary cluster-size inference with random field and permutation methods. Neuroimage. 2004;22:676–687.

44. Worsley KJ, Taylor JE, Tomaiuolo F, Lerch J. Unified univariate and multivariate random field theory. NeuroImage. 2004;23:S189–S195.

45. Herculano-Houzel S, Watson CR, Paxinos G. Distribution of neurons in functional areas of the mouse cerebral cortex reveals quantitatively different cortical zones. Front Neuroanat. 2013;7.

46. Ashburner M, Ball CA, Blake JA, Botstein D, Butler H, Cherry JM, et al. Gene Ontology: tool for the unification of biology. Nature Genetics. 2000;25:25–29.

47. The Gene Ontology Consortium. The Gene Ontology Resource: 20 years and still GOing strong. Nucleic Acids Res. 2019;47:D330–D338.

48. Kang HJ, Kawasawa YI, Cheng F, Zhu Y, Xu X, Li M, et al. Spatio-temporal transcriptome of the human brain. Nature. 2011;478:483–489.

49. Paquola C, Seidlitz J, Benkarim O, Royer J, Klimes P, Bethlehem RAI, et al. A multi-scale cortical wiring space links cellular architecture and functional dynamics in the human brain. PLOS Biology. 2020;18:e3000979.

50. von Economo CF, Koskinas GN. Die cytoarchitektonik der hirnrinde des erwachsenen menschen. J. Springer; 1925.

51. Scholtens LH, de Reus MA, de Lange SC, Schmidt R, van den Heuvel MP. An MRI Von Economo - Koskinas atlas. Neuroimage. 2018;170:249–256.

52. Alexander-Bloch AF, Shou H, Liu S, Satterthwaite TD, Glahn DC, Shinohara RT, et al. On testing for spatial correspondence between maps of human brain structure and function. Neuroimage. 2018;178:540–551.

53. Burt JB, Helmer M, Shinn M, Anticevic A, Murray JD. Generative modeling of brain maps with spatial autocorrelation. NeuroImage. 2020;220:117038.

54. Markello RD, Misic B. Comparing spatially-constrained null models for parcellated brain maps. BioRxiv. 2020:2020.08.13.249797.

55. Desikan RS, Ségonne F, Fischl B, Quinn BT, Dickerson BC, Blacker D, et al. An automated labeling system for subdividing the human cerebral cortex on MRI scans into gyral based regions of interest. NeuroImage. 2006;31:968–980.

56. Schmaal L, Hibar DP, Sämann PG, Hall GB, Baune BT, Jahanshad N, et al. Cortical abnormalities in adults and adolescents with major depression based on brain scans from 20 cohorts worldwide in the ENIGMA Major Depressive Disorder Working Group. Mol Psychiatry. 2017;22:900–909.

57. Hibar DP, Westlye LT, Doan NT, Jahanshad N, Cheung JW, Ching CRK, et al. Cortical abnormalities in bipolar disorder: an MRI analysis of 6503 individuals from the ENIGMA Bipolar Disorder Working Group. Molecular Psychiatry. 2018;23:932–942.

58. Hoogman M, Muetzel R, Guimaraes JP, Shumskaya E, Mennes M, Zwiers MP, et al. Brain Imaging of the Cortex in ADHD: A Coordinated Analysis of Large-Scale Clinical and Population-Based Samples. AJP. 2019;176:531–542.

59. Boedhoe PSW, Schmaal L, Abe Y, Alonso P, Ameis SH, Anticevic A, et al. Cortical Abnormalities Associated With Pediatric and Adult Obsessive-Compulsive Disorder: Findings From the ENIGMA Obsessive-Compulsive Disorder Working Group. AJP. 2017;175:453–462.

60. van Rooij D, Anagnostou E, Arango C, Auzias G, Behrmann M, Busatto GF, et al. Cortical and Subcortical Brain Morphometry Differences Between Patients With Autism Spectrum Disorder and Healthy Individuals Across the Lifespan: Results From the ENIGMA ASD Working Group. Am J Psychiatry. 2018;175:359–369.

61. Larivière S, Paquola C, Park B, Royer J, Wang Y, Benkarim O, et al. The ENIGMA Toolbox: multiscale neural contextualization of multisite neuroimaging datasets. Nat Methods. 2021;18:698–700.

62. Sawyer SM, Azzopardi PS, Wickremarathne D, Patton GC. The age of adolescence. Lancet Child Adolesc Health. 2018;2:223–228.

63. Meng X, Rosenthal R, Rubin DB. Comparing correlated correlation coefficients. Psychological Bulletin. 1992;111:172–175.

64. Collins CE, Airey DC, Young NA, Leitch DB, Kaas JH. Neuron densities vary across and within cortical areas in primates. Proc Natl Acad Sci U S A. 2010;107:15927–15932.

65. Cahalane DJ, Charvet CJ, Finlay BL. Systematic, balancing gradients in neuron density and number across the primate isocortex. Front Neuroanat. 2012;6.

66. Carlo CN, Stevens CF. Structural uniformity of neocortex, revisited. PNAS. 2013;110:1488–1493.

67. Braitenberg V, Schüz A. Density of Axons. In: Braitenberg V, Schüz A, editors. Cortex: Statistics and Geometry of Neuronal Connectivity, Berlin, Heidelberg: Springer Berlin Heidelberg; 1998. p. 39–42.

68. Economo C von. Cellular Structure of the Human Cerebral Cortex. Karger; 2009.

69. Berdenis van Berlekom A, Muflihah CH, Snijders GJLJ, MacGillavry HD, Middeldorp J, Hol EM, et al. Synapse Pathology in Schizophrenia: A Meta-analysis of Postsynaptic Elements in Postmortem Brain Studies. Schizophrenia Bulletin. 2020;46:374–386.

70. Srinivas KV, Buss EW, Sun Q, Santoro B, Takahashi H, Nicholson DA, et al. The Dendrites of CA2 and CA1 Pyramidal Neurons Differentially Regulate Information Flow in the Cortico-Hippocampal Circuit. J Neurosci. 2017;37:3276–3293.

71. Cannon TD, Chung Y, He G, Sun D, Jacobson A, van Erp TGM, et al. Progressive Reduction in Cortical Thickness as Psychosis Develops: A Multisite Longitudinal Neuroimaging Study of Youth at Elevated Clinical Risk. Biological Psychiatry. 2015;77:147–157.

72. Consortium TB, Anttila V, Bulik-Sullivan B, Finucane HK, Walters RK, Bras J, et al. Analysis of shared heritability in common disorders of the brain. Science. 2018;360.

73. Kirov G, Pocklington AJ, Holmans P, Ivanov D, Ikeda M, Ruderfer D, et al. De novo CNV analysis implicates specific abnormalities of postsynaptic signalling complexes in the pathogenesis of schizophrenia. Molecular Psychiatry. 2012;17:142–153.

74. Consortium TSWG of the PG, Ripke S, Walters JT, O’Donovan MC. Mapping genomic loci prioritises genes and implicates synaptic biology in schizophrenia. MedRxiv. 2020:2020.09.12.20192922.

75. Ball G, Beare R, Seal ML. Charting shared developmental trajectories of cortical thickness and structural connectivity in childhood and adolescence. Hum Brain Mapp. 2019;40:4630–4644.

76. Sotiras A, Toledo JB, Gur RE, Gur RC, Satterthwaite TD, Davatzikos C. Patterns of coordinated cortical remodeling during adolescence and their associations with functional specialization and evolutionary expansion. PNAS. 2017;114:3527–3532.

77. Zielinski BA, Prigge MBD, Nielsen JA, Froehlich AL, Abildskov TJ, Anderson JS, et al. Longitudinal changes in cortical thickness in autism and typical development. Brain. 2014;137:1799–1812.

78. Di Martino A, Fair DA, Kelly C, Satterthwaite TD, Castellanos FX, Thomason ME, et al. Unraveling the Miswired Connectome: A Developmental Perspective. Neuron. 2014;83:1335–1353.

79. Klingler E, Francis F, Jabaudon D, Cappello S. Mapping the molecular and cellular complexity of cortical malformations. Science. 2021;371.

